# Cholangiocarcinoma presents a distinct myeloid-derived suppressor cell signature compared to other hepatobiliary cancers

**DOI:** 10.1101/554600

**Authors:** Defne Bayik, Adam J. Lauko, Gustavo A. Roversi, Emily Serbinowski, Lou-Anne Acevedo-Moreno, Christopher Lanigan, Mushfig Orujov, Alice Lo, Tyler J. Alban, Daniel J. Silver, J. Mark Brown, Daniela S. Allende, Federico N. Aucejo, Justin D. Lathia

## Abstract

Myeloid-derived suppressor cells (MDSCs) are immunosuppressive cells that are increased in patients with numerous malignancies including viral-derived hepatocellular carcinoma (HCC). Here, we report an elevation of MDSC in other hepatobiliary malignancies including non-viral HCC, neuroendocrine tumors (NET), colorectal carcinoma with liver metastases (CRLM), but not cholangiocarcinoma (CCA). Investigation of myeloid cell infiltration in HCC, NET and intrahepatic CCA tumors further established that the frequency of antigen-presenting cells was limited compared to benign lesions suggesting that primary and metastatic hepatobiliary cancers have distinct peripheral and tumoral myeloid signatures. Bioinformatics analysis of the Cancer Genome Atlas demonstrated that a high MDSC score in HCC patients predicted poor disease outcome. Mechanistic studies indicated that the oncometabolite D-2-hydroxyglutarate resulting from isocitrate dehydrogenase 1 mutation could be a limiting factor of MDSC accumulation in CCA patients. Given our observation that MDSCs are increased in non-CCA malignant liver cancers, they may comprise suitable targets for effective immunotherapy approaches.

## Introduction

Primary hepatocellular carcinoma (HCC) is among the leading cause of cancer-related deaths in the U.S., with 40,000 new cases and 30,000 mortalities in 2018^1^. Chronic hepatitis B virus (HBV) and hepatitis C virus (HCV) infection, obesity, and excess alcohol consumption are risk factors associated with HCC^2^. HCC resulting from chronic viral infections constitute 75% of all cases, while the non-viral causes account for the remaining 25%. Secondary liver cancers that originate by the metastatic spread of a primary tumor from a distant site, most commonly colorectal carcinoma (CRLM), are more frequent than primary HCC^3^. Cholangiocarcinoma (CCA) and neuroendocrine tumors (NETs) are rare hepatobiliary cancers, which have unique disease presentation compared to HCC and CRLM, and distinct therapeutic responsiveness^4,5^. Current treatment strategies including surgical resection, radiation, ablation, embolization, systemic/local chemotherapy infusion and liver transplantation have improved the outcome of patients with HCC and CCA, however, liver cancers are diagnosed at advanced stages with poor clinical presentation and have high recurrence rates^6^. Thus, there is a need to gain mechanistic insight into the pathobiology of liver cancers and develop more effective therapeutic opportunities.

Tumors employ multiple mechanisms to evade immune recognition. Myeloid-derived suppressor cells (MDSCs) are a heterogenous population of immature myeloid cells that expand in patients with malignancies and infiltrate tumors, where they confine anti-tumor immune response by suppressing cytotoxic T cell and natural killer (NK) cell activity^7^. Previous work demonstrated that high MDSC frequency in melanoma not only correlates with poor patient outcome but also associates with therapeutic irresponsiveness^7-10^. Targeting MDSCs in multiple preclinical models including colorectal and liver cancers activated an anti-tumor immune response and reduced tumor growth, suggesting that modulating these cells is a promising strategy for cancer immunotherapy^11,12^. Several studies have also reported that MDSCs are augmented in peripheral blood of HCC patients^13–16^. However, the relevance of this observation in terms of other hepatobiliary malignancies, including non-viral HCC, and patient clinical presentation has yet to be determined. Considering the range and pathobiological variation in primary and metastatic liver cancers, we hypothesized that MDSCs might have distinct associations with different types of hepatobiliary tumors. By screening blood and liver surgical specimens from patients with HCC, CCA, CRLM and NET, we identified that MDSCs are elevated in most but not all liver cancers. Specifically, MDSC levels remained low in patients with CCA, potentially due to the inhibitory function of the oncometabolite D-2-Hydroxyglutarate (D2HG) produced by the mutant isocitrate dehydrogenase (IDH)1/2 enzyme. Our findings demonstrate that MDSC augmentation is impacted by the type of liver tumors, and support testing of MDSC targeting strategies in patients with HCC, CRLM and NET but not CCA cases.

## Materials and Methods

### Reagents

Fluorophore-conjugated anti-human CD3 (UCHT1), CD4 (SK3), CD8 (RPA-T8), CD14 (MφP9), CD15 (HI98), CD25 (M-A251), CD33 (WM53), CD107a (H4A3), CD127 (HIL-7R-M21) and HLA-DR (G46-6) antibodies were purchased from BD Biosciences. Anti-mouse Ly6G (1A8), Ly6C (HK1.4), CD11b (M1/70) and CD3 (145-2C11) were purchased from Biolegend. Anti-human CD11b (CD11B29) antibody, anti-mouse Gr-1 (RB6-8C5) antibody and LIVE/DEAD™ Fixable Blue Dead Cell Stain Kit, for UV excitation was obtained from ThermoFisher Scientific.

### Patients

Peripheral blood samples were collected perioperatively from 114 patients with non-viral liver cancer in the Cleveland Clinic. All patients provided written informed consent under IRB 10-347 and the protocol was approved by the Cleveland Clinic Institutional Review Board. All the diagnoses were confirmed by the Cleveland Clinic Pathology Department. Patient clinical characteristics are provided in **Table 1**.

**Table 1:**
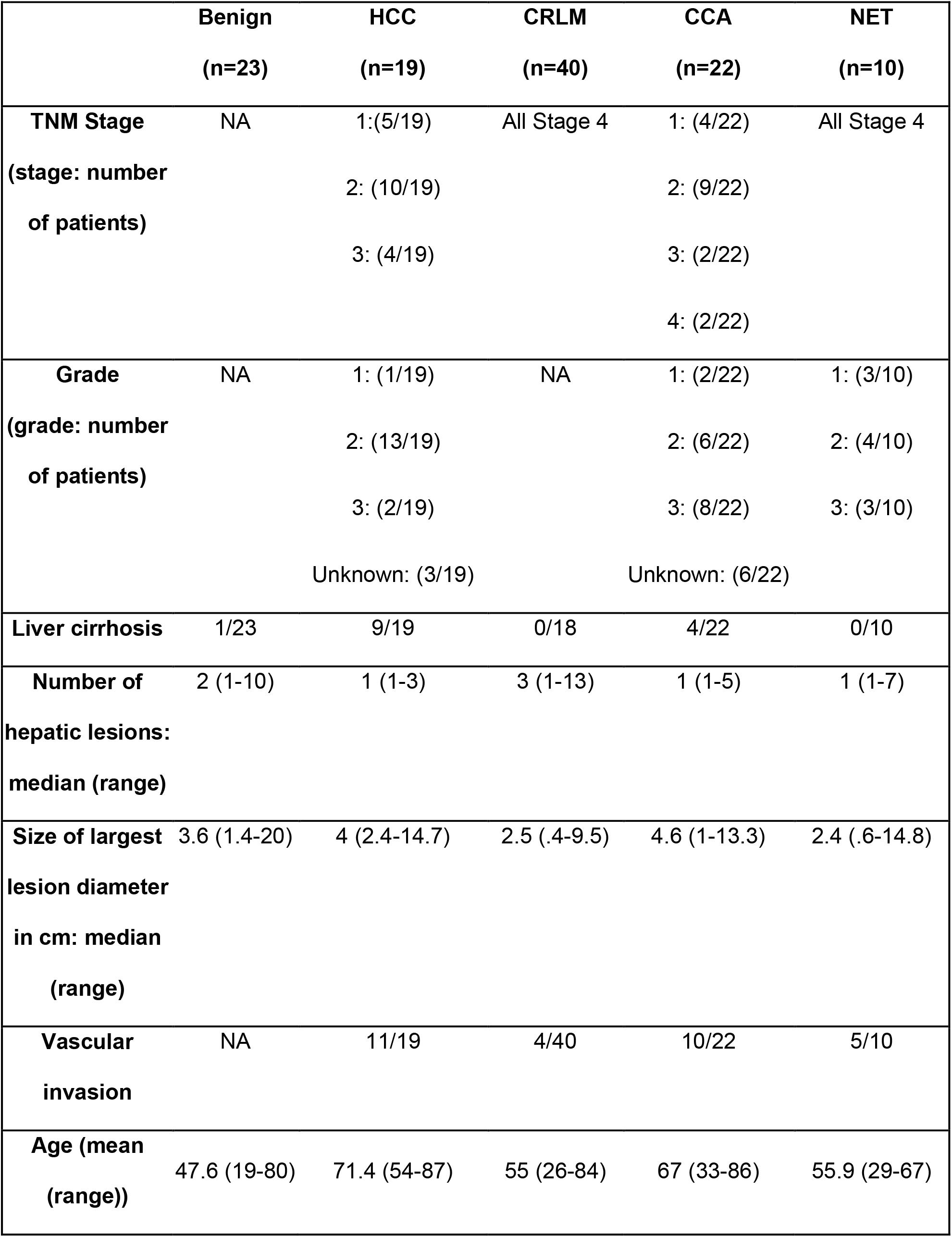
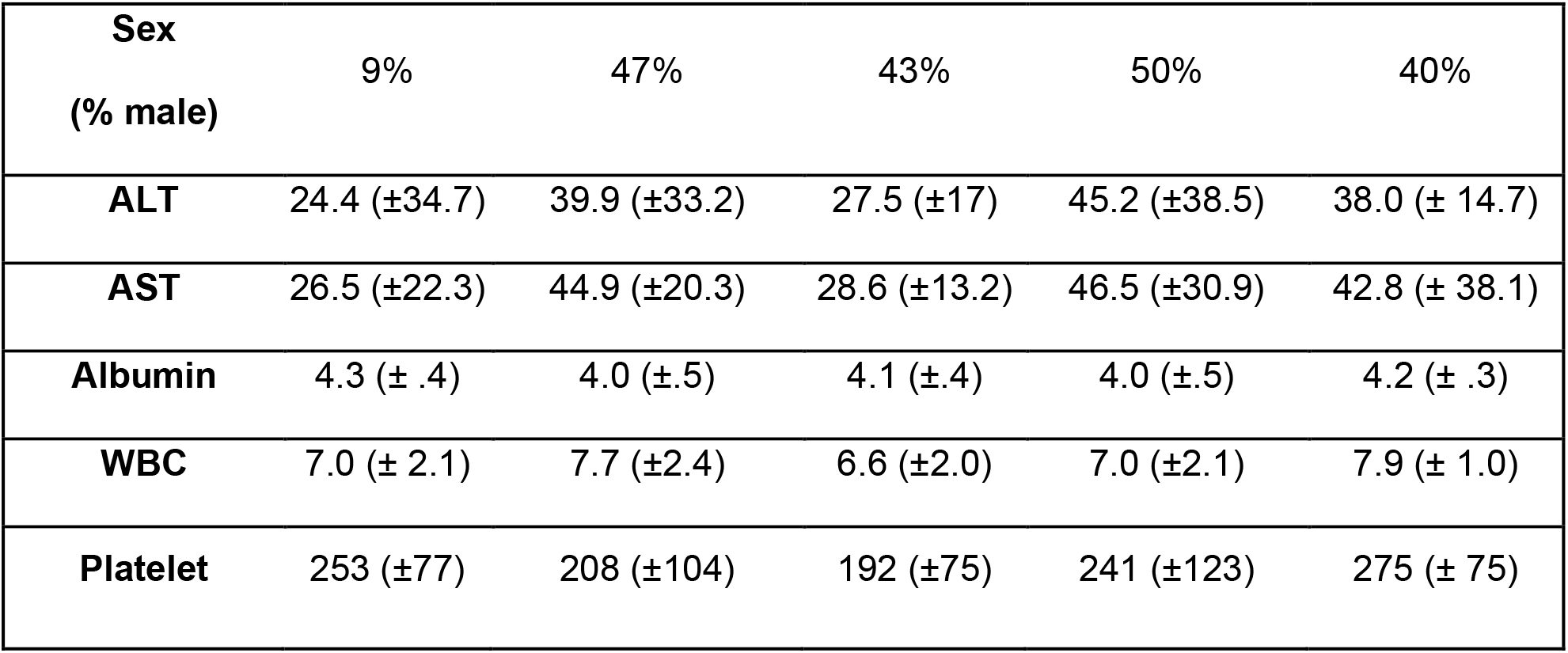
Patient demographic information

|  | **Benign**<br>**(n=23)** | **HCC**<br>**(n=19)** | **CRLM**<br>**(n=40)** | **CCA**<br>**(n=22)** | **NET**<br>**(n=10)** |
| --- | --- | --- | --- | --- | --- |
| **TNM Stage**<br>(stage: number<br>of patients) | NA | 1:(5/19)<br>2: (10/19)<br>3: (4/19) | All Stage 4 | 1: (4/22)<br>2: (9/22)<br>3: (2/22)<br>4: (2/22) | All Stage 4 |
| **Grade**<br>(grade: number<br>of patients) | NA | 1: (1/19)<br>2: (13/19)<br>3: (2/19)<br>Unknown: (3/19) | NA | 1: (2/22)<br>2: (6/22)<br>3: (8/22)<br>Unknown: (6/22) | 1: (3/10)<br>2: (4/10)<br>3: (3/10) |
| **Liver cirrhosis** | 1/23 | 9/19 | 0/18 | 4/22 | 0/10 |
| **Number of**<br>**hepatic lesions:**<br>**median (range)** | 2 (1-10) | 1 (1-3) | 3 (1-13) | 1 (1-5) | 1 (1-7) |
| **Size of largest**<br>**lesion diameter**<br>**in cm: median**<br>**(range)** | 3.6 (1.4-20) | 4 (2.4-14.7) | 2.5 (.4-9.5) | 4.6 (1-13.3) | 2.4 (.6-14.8) |
| **Vascular**<br>**invasion** | NA | 11/19 | 4/40 | 10/22 | 5/10 |
| **Age (mean**<br>**(range))** | 47.6 (19-80) | 71.4 (54-87) | 55 (26-84) | 67 (33-86) | 55.9 (29-67) |

| **Sex**<br>**(% male)** | 9% | 47% | 43% | 50% | 40% |
| --- | --- | --- | --- | --- | --- |
| **ALT** | 24.4 (±34.7) | 39.9 (±33.2) | 27.5 (±17) | 45.2 (±38.5) | 38.0 (± 14.7) |
| **AST** | 26.5 (±22.3) | 44.9 (±20.3) | 28.6 (±13.2) | 46.5 (±30.9) | 42.8 (± 38.1) |
| **Albumin** | 4.3 (± .4) | 4.0 (±.5) | 4.1 (±.4) | 4.0 (±.5) | 4.2 (± .3) |
| **WBC** | 7.0 (± 2.1) | 7.7 (±2.4) | 6.6 (±2.0) | 7.0 (±2.1) | 7.9 (± 1.0) |
| **Platelet** | 253 (±77) | 208 (±104) | 192 (±75) | 241 (±123) | 275 (± 75) |

### Quantification of MDSC in peripheral blood

Peripheral blood samples were processed using a Ficoll-Paque PLUS gradient in a SepMateTM (Stem Cell Technologies) and frozen in 10% DMSO (Fisher Scientific) and 90% FBS (Gibco) until further analysis. Samples were stained with live/dead UV stain (Invitrogen) and blocked in FACS buffer (PBS, 2% BSA) containing an FcR blocking reagent (Miltenyi Biotec). Samples were stained with fluorophore-conjugated antibody cocktails, washed with FACS buffer and analyzed using LSRFortessa (BD Biosciences). Analysis were performed using FlowJo software (Tree Star Inc.).

### Immunohistochemistry (IHC)

Formalin fixed paraffin-embedded blocks and H&E stained slides from liver biopsy specimens were available in 29 patients (6 HCC, 9 benign, 10 CCA and 4 NET) for IHC analysis. IHC was performed using the mouse monoclonal CD33 (PWS44, Leica Biosystems) and HLA-DR/DP/DQ (CR3/43, ThermoFisher) antibody diluted 1:100 using SignalStain Diluent (Cell Signaling Technology) and incubated for 40 minutes at 37°C. Automated staining used the VENTANA BenchMark Ultra platform, U iView DAB software, and was detected using iView DAB with the Endogenous Biotin Blocking Kit (Ventana Medical Systems Inc.). Antigen Retrieval used Cell Conditioning#1 selected for the Standard Time. Normal lymph node tissue was used for positive run controls. For CD68 staining, samples were incubated with pre-diluted CD68 antibody (clone KP-1, Ventana) for 16 minutes at 37°C and analyzed with OptiView DAB detection kit. IHC staining conditions are summarized in **Supplemental Figure 1**.

HLA-DR positive cells were assessed quantitatively within the lesional tissue, at a 200X magnification on an Olympus B41 microscope. Similarly, CD33 and CD68 positive cells were counted at the periphery of the lesion (tumor-normal interface) and intralesionally at 200x. For all three markers, the peak count was considered.

### Bioinformatic Analysis of Immune Populations

The Cancer Genome Atlas (TCGA) Liver Cancer database was accessed via University of California Santa Cruz (UCSC) Xena Browser [https://xenabrowser.net/heatmap/] on March 21, 2018 and December 13, 2018. CD33, CD11b (ITGAM), HLA-DRA, CD4, CD25 (IL-2RA), FoxP3 and CD8a expression levels, patient demographic information, viral status and overall survival data were extracted for further analysis. To generate MDSC signature, only samples with CD33 and CD11b expression above the median values were used. From this set, samples with low HLA-DRA expression were categorized into “high MDSC signature” group. CD33^high^CD11b^high^HLA-DRA^high^ samples were used for the control population. Treg signature was generated based on initial CD4 positive gating and subsequent high CD25 and FoxP3 expression levels. Samples with CD25 and FoxP3 levels below the median served as controls. CD8a^high^CD107a^high^ samples were considered to have high levels of activated CD8 T cells, while samples with CD8a^high^CD107a^low^ expression levels were classified as controls.

### Bone marrow-derived MDSCs

MDSCs were polarized by stimulating bone marrows of 8-10 weeks of male C57BL/6 mice with 40ng/ml GM-CSF and 80ng/ml IL-13 for 3-4 days^17^. Bone marrow cells were treated with 0.3 mM-30 mM D-2-Hydroxyglutarate (D2HG, Cayman Chemical) and Alpha-ketoglutarate (αKG, Sigma Aldrich) during the polarization step. MDSC phenotype was confirmed by staining the cells in FACS buffer with 1:100 diluted anti-CD11b, Ly6C and Ly6G antibodies after live/dead dye staining and FcR blocking. Samples were acquired with LSRFortessa and analyzed using FlowJo software.

For functional confirmation, MDSC and nonMDSC fraction were sorted using biotin-conjugated anti-Gr-1 antibody and streptavidin microbeads (Miltenyi Biotec) based on manufacturer’s instructions. Increased ratios of MDSCs and nonMDSCs were co-cultured with splenic T cells isolated with Pan T Cell Isolation Kit II (Miltenyi Biotec) and stained with 1 μM Carboxyfluorescein succinimidyl ester (CFSE, Biolegend). Stimulation of T cells was achieved by adding 40 IU IL-2 and Dynabeads Mouse T-Activator CD3/CD28 (ThermoFisher Scientific) for 4 days. Samples were stained with anti-CD11b and anti-CD3 at the end of the incubation period and analyzed using LSRFortessa. Proliferation rate of CD3^+^CD11b^-^ cells was determined with FlowJo.

### Statistical Analysis

Two-way ANOVA or Student’s t-test was utilized to test differences in continuous variables with normal distribution. Kruskal-Wallis test was performed as nonparametric test when normal distribution could not be assumed. The long-rank test was utilized to determine difference in overall survival. p-value <0.05 was considered statistically significant (GraphPad Software Inc., JMP Pro and Version 14. SAS Institute Inc.).

## Results

### Patient Characteristics

To gain a more comprehensive understanding of the link between MDSCs and liver cancers, we analyzed MDSC frequency from peripheral blood mononuclear cells (PBMC) of 114 patients with diagnoses ranging from benign lesions, HCC, CRLM, NET to CCA. Characteristics of these patients are summarized in **Table 1**. Briefly, 19 patients were diagnosed with HCC, grade 1-3 and TNM stage 1-3. Of the 9 HCC patients with cirrhosis, only one was a Class B Child-Pugh Score (remainder Class A). The benign lesions were primarily hepatic adenomas, with two adrenocortical adenoma, two focal nodular hyperplasia, one biliary cystadenoma, three hemangioma, two biliary cystadenoma, one pseudolipoma of Glisson’s capsule, one fibrotic cyst and one patient with polycystic liver disease. No CRLM patients had cirrhosis and intrahepatic lesion number ranged from 1-13. The CCA patient’s tumors ranged from TNM stage 1-4 and grade 1-3 and all but one patient had intrahepatic CCA. Ten out of twenty-two CCA patients had vascular invasion. The NET ranged from grade 1-3 and included primary tumors in the following locations: small bowel (2), terminal ileum, pancreas, stomach, adrenocortical (2), large bowel and unknown primary (2). HCC and CCA patients compared to patients with benign tumors were older (median age 71.4/67.5/47.6, p<0.0001) and more likely to be male (p<0.01). HCC patients also presented elevated levels of liver enzymes (ALT 39.9 vs 24.4, p=0.2 and AST 44.9 vs 23.4, p=0.01), a trend also observed in CCA patients (ALT 45.2 vs 24.4, p=0.06 and AST 46.5 vs 26.5, p<0.05).

### MDSCs are increased in peripheral blood of primary and metastatic liver tumor patients

MDSCs are bone marrow-derived immature immunosuppressive cells that accumulate in patients with malignancies^9,18^. In human, MDSCs are broadly categorized into monocytic (mMDSCs, CD33^+^CD11b^+^HLA-DR^-/low^CD14^+^) and granulocytic (gMDSC, CD33^+^CD11b^+^HLA-DR^-/low^CD14^-^) subsets, which differ in their suppressive capacity^7,19^. Previous studies demonstrated that the monocytic subset of MDSCs are increased in circulation of HCC patients and their levels are impacted by radiation therapy^13–16^. As previously reported, CD33^+^CD11b^+^HLA-DR^-/low^ MDSC frequency was elevated in HCC patients compared to patient with benign liver lesions, and showed a trend towards increased abundance with tumor grade (**Fig 1B, Supplemental Fig 1A**). Similarly, patients with NET and CRLM had a significant increase in the circulating MDSC frequency (**Fig 1B**). In contrast, MDSC percentages in CCA patients were indistinguishable from those with benign liver lesions independent of tumor grade (**Fig 1B, Supplemental Fig 1B**). Prior studies suggest that mMDSCs are the main population of MDSCs in patients with liver cancers^14–16^. Consistently, our findings demonstrated that CD14^+^CD15^-^ mMDSC subset constitute the majority of MDSCs in peripheral circulation, and that these cells are present at higher rates in patients with HCC compared to benign lesions (**Fig 1C-D**). Regulatory T cell (Tregs) constitute another major immunosuppressive cell population frequently increases in patients with cancer that suppresses activity of anti-tumoral CD8^+^ T cells in the tumor microenvironment^20^. It has been established that MDSCs and Tregs cross-talk to induce the generation and recruitment of one another^21^. We analyzed the frequency of Tregs and activated CD8^+^ T cells to investigate whether these populations correlate with disease status. Neither Tregs nor activated CD8^+^ T cells were significantly different among various types of primary and metastatic liver malignancies (**Fig 1E-F**). Together, these results indicate that MDSCs are linked to malignancy but CCA represent an exception with lower MDSC frequency.

**Figure 1.**
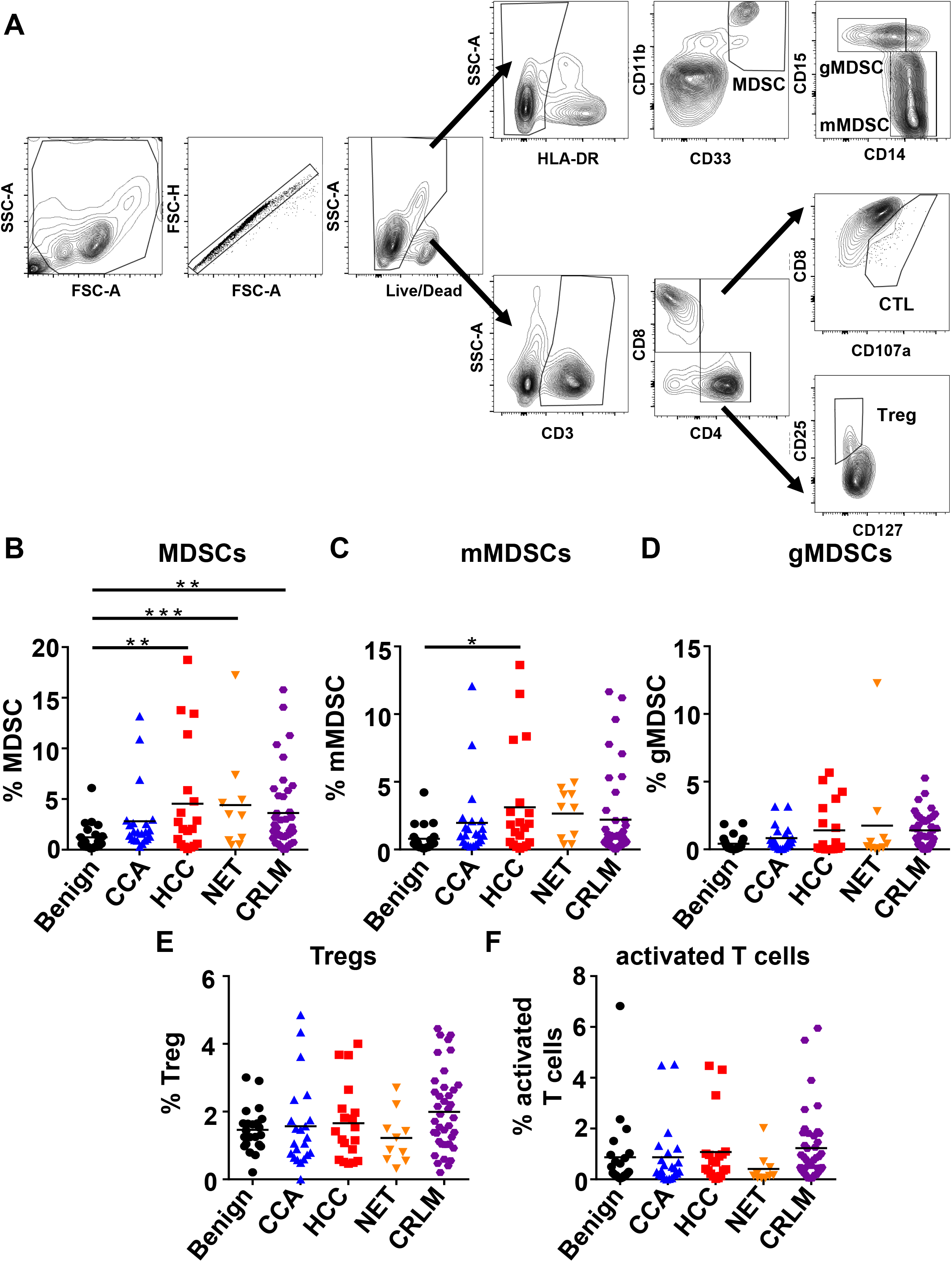
Frequency of MDSCs is increased in HCC, CRLM and NET patients. A) Representative dot plots depicting the gating strategy of different immune populations. Frequency of B) CD33^+^CD11b^+^HLA-DR^-/low^ MDSC, C) CD33^+^CD11b^+^HLA-DR^-/low^CD14^+^ mMDSC, D) CD33^+^CD11b^+^HLA-DR^-/low^CD14^-^ gMDSC, E) CD3^+^CD4^+^CD25^+^CD127^-^ Treg, and F) CD3^+^CD8^+^CD107a^+^ activated T cells in PBMC of CRLM (n=41), HCC (n=18), CCA (n=18), NET (n=5) and benign (n=18) cases were analyzed with flow cytometry. * p<.05; ** p<0.01; *** p<0.001 as determined by 2-way ANOVA.

To investigate whether MDSC frequency correlates with clinical parameters, our patient cohort was split into a MDSC high (n=23) and MDSC low (n=86) group based on the mean peripheral MDSC percentage (4.5%) in the HCC subgroup. There was no significant difference between AST (38.3. vs 34.7, p=0.5) and ALT (36.6 vs 32.6, p=0.6) between MDSC high versus low groups (**Supplemental Fig 2A**).To examine whether MDSC frequency is linked to tumor markers we investigated HCC and CRLM., There was no statistical correlation between MDSC frequency and AFP levels in patients with HCC (p=.09), but high MDSC levels (>4.5%) correlated with elevated CEA in CRLM patients (**Supplemental Fig 2B-C**).

### Benign liver lesions are infiltrated by antigen-presenting cells (APCs)

Under normal physiological conditions, mMDSCs are capable of differentiating into antigen-presenting dendritic cells (DCs) and macrophages^7^. This pathway has been hypothesized to be halted under pathophysiological conditions such as cancer, leading to accumulation of these cells in host. Thus, we investigated whether enhanced MDSC frequency associated with APC infiltration of liver lesions by performing immunohistochemistry staining of pan-myeloid marker CD33, MHC Class II (HLA-DR) and macrophage marker CD68 in a subset of HCC, CCA, NET and benign cases. Malignant tumors exhibited a similar immune profile with no significant differences in the number of CD33, HLA-DR and CD68 positive cells in peak areas counted within the tumors (**Fig 2A-B**). Compared to HCC and CCA, benign lesions had higher numbers of intralesional HLA-DR or CD33 positive cells (HLA-DR^+^ cells in HCC-CCA 36.5 vs 54.9 in benign lesions, p<0.05; CD33^+^ cells in HCC-CCA 16.7 vs 39.2 in benign lesions, p<0.05; **Fig 2C**), representing APCs. CD68^+^ cells corresponding to macrophages, including resident Kupffer cells, were not different between malignant and benign liver lesions (**Fig 2B-C**), suggesting that DCs constitute the majority of infiltrating APCs in benign cases. In addition, we analyzed whether the immunosuppressive tumor microenvironment prevents infiltration of myeloid cells leading to the accumulation of these cells at the periphery (tumor-normal interface). Intralesional versus peripheral specimens from the matching tumors had similar numbers of CD33 or CD68 positive cells, CCA cases being the exception (**Fig 2D**). These results suggest that HCC and NETs result in reduced myeloid cell accumulation in the tumor as well as the adjacent tissue.

**Figure 2.**
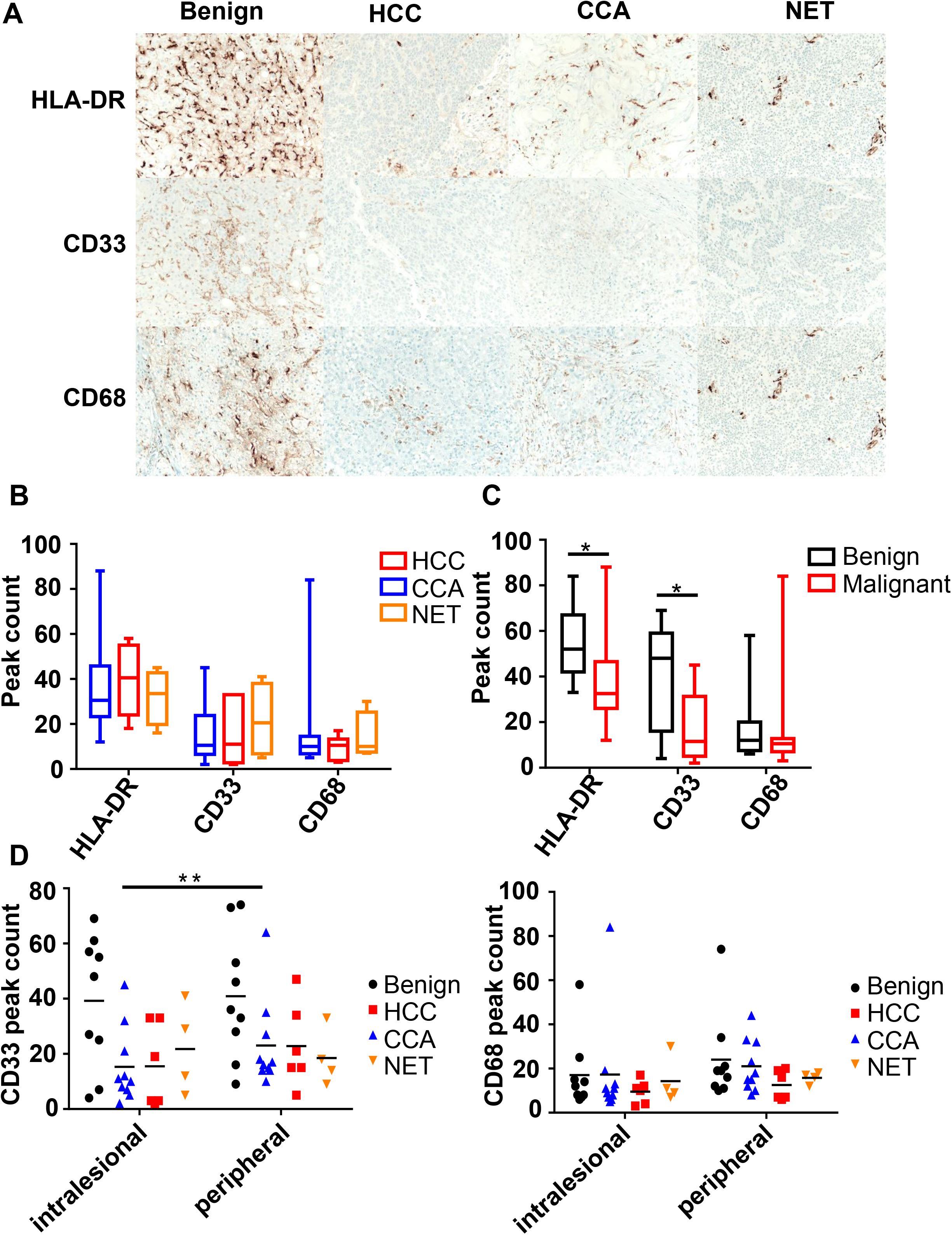
HCC, NET and CCA tumors have similar myeloid cell infiltration pattern. A) Representative immunohistochemistry images demonstrating infiltration HLA-DR^+^ APCs, CD33^+^ myeloid cells and CD68^+^ macrophages. Quantification of peak area count of intralesional HLA-DR, CD33 and CD68 staining B) from HCC (n=6), CCA (n=10) and NET (n=4) cases C) from malignant liver cancers (n=20) versus benign lesions (n=9). D) Comparison of intralesional versus peripheral CD33 and CD68 peak area count. Data represented as min-to-max and *<p=0.05 as determined individually for each marker by two-tailed t-test with unequal variance; **<p=0.01 as by paired two-tailed t-test.

### MDSC frequency correlates with HCC outcome

Although HCC patients had significantly higher levels of MDSCs compared to patients with benign lesions, this patient subpopulation clustered into two groups based on MDSC levels in PBMCs (MDSC high (>4.5%) and low (<4.5%)) (**Fig 1B**). Based on this variation within the PBMCs, we investigated whether MDSC frequency within the tumor correlates with disease outcome using the TCGA dataset containing gene expression information from 440 HCC patients. We generated a high versus low MDSC score by focusing on high CD33 and CD11b expression and further creating subgroups based on the median levels of HLA-DR expression. Patients with a high MDSC score (CD33^high^CD11b^high^HLA-DR^low^) had significantly reduced overall survival duration, compared to those with low MDSC score (CD33^high^CD11b^high^HLA-DR^high^; **Fig 3A**). Importantly, these survival differences were not dependent on tumor grade, viral status or biological sex as no significant difference for these parameters was detected between cohorts (*data not shown*). Using the same strategy, we generated signatures for immunosuppressive Tregs (CD4^high^CD25^high^FoxP3^high^) and activated CD8^+^ T cells (CD8^high^CD107a^high^). There was no survival difference in patients with a high Treg signature compared to those with a low Treg (CD4^high^) score (**Fig 3B**). Similarly, an activated CD8^+^ T cell signature did not predict survival advantage over having more naïve CD107a^low^ T cells in the tumor microenvironment (**Fig 3C**). Together, these observations suggest that MDSCs but not Tregs or activated CD8^+^ T cells associate with HCC outcome.

**Figure 3.**
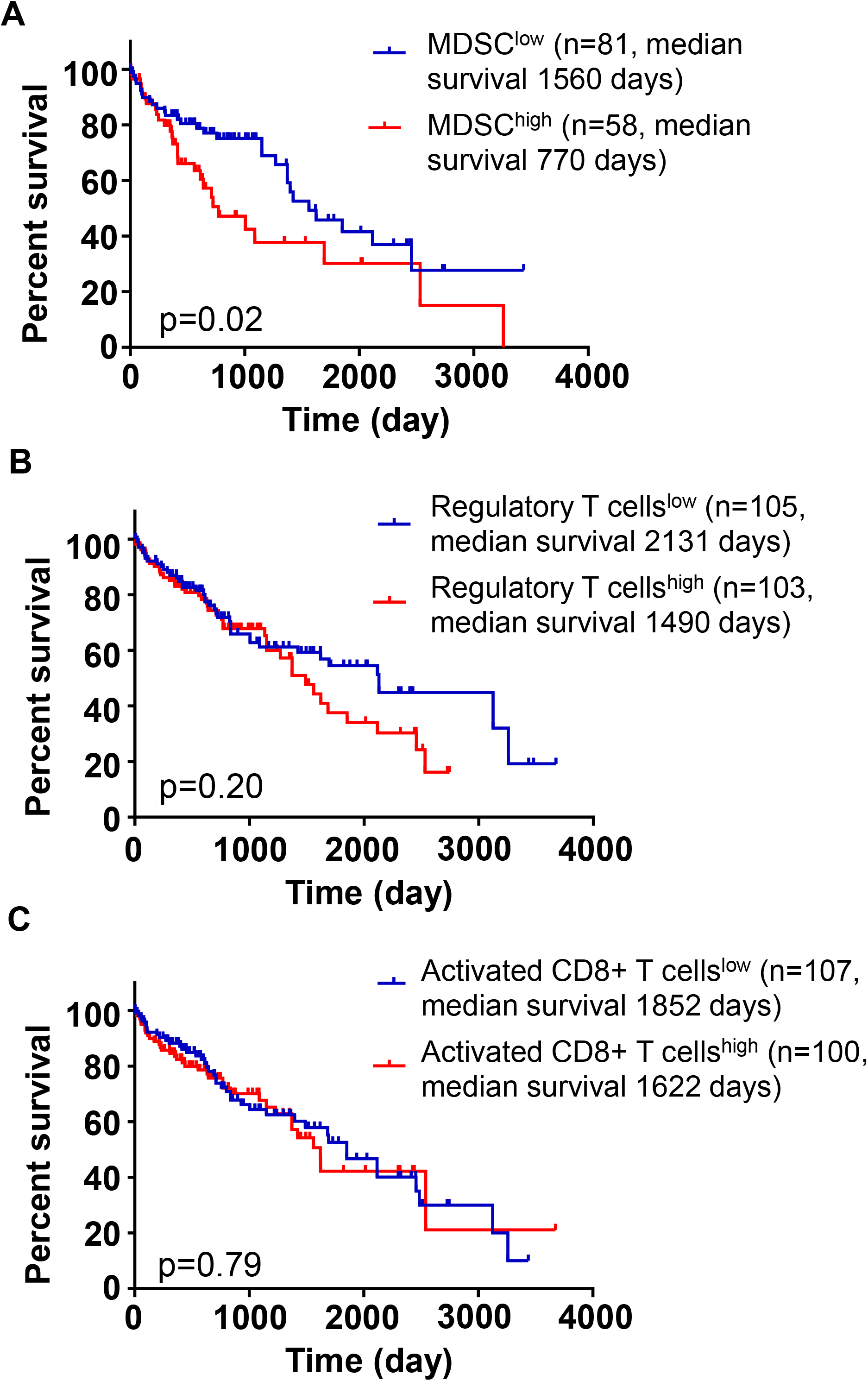
High MDSC score predicts poor HCC prognosis. Kaplan-Meier curved depicting overall survival of HCC patients with high A) MDSC (CD33^+^CD11b^+^HLA-DR^-/low^), B) Treg (CD4^+^CD25^+^FoxP3^+^) and C) Activated T cell (CD8a^+^CD107a^+^) score. Data are from TCGA and were obtained from UCSC Xena Browser. p <0.05 was determined by Log-rank (Mantel-Cox) test.

### MDSC frequency is regulated by D2HG

CCA bear a different set of genomic alterations compared to HCC, and 25% of the cases have reported to have IDH1 mutation^22^. Mutant IDH1 enzyme converts α-ketoglutarate (αKG), a byproduct of Krebs cycle and a common substrate of histone modifying enzymes, to the oncometabolite D2HG^23^. Prior studies in glioma have suggested that IDH mutations can alter the immune profile and affect T cell response, therefore we analyzed the role of D2HG in MDSC regulation. We polarized MDSCs from bone marrow cells by stimulating with GM-CSF and IL-13 and confirmed MDSCs phenotypically based on flow staining (**Fig 4A**). Functionality of MDSCs was further verified by their ability to suppress the proliferation of activated T cells (**Fig 4B**). To test the differential effects of αKG versus D2HG, we included these metabolites during the MDSC polarization period. While αKG had no significant impact on the generation of MDSCs, D2HG significantly reduced the frequency of mMDSCs in the cultures without interfering with gMDSC polarization (**Fig 4C**). Combined with the observation that mMDSCs is the prevalent subset in the circulation of patients with hepatobiliary malignancies with the exception of CCA cases and that D2HG levels are elevated in the serum of CCA patients^24^, these results point to a potential mechanism of MDSC suppression via D2HG accumulation in CCA patients.

**Figure 4.**
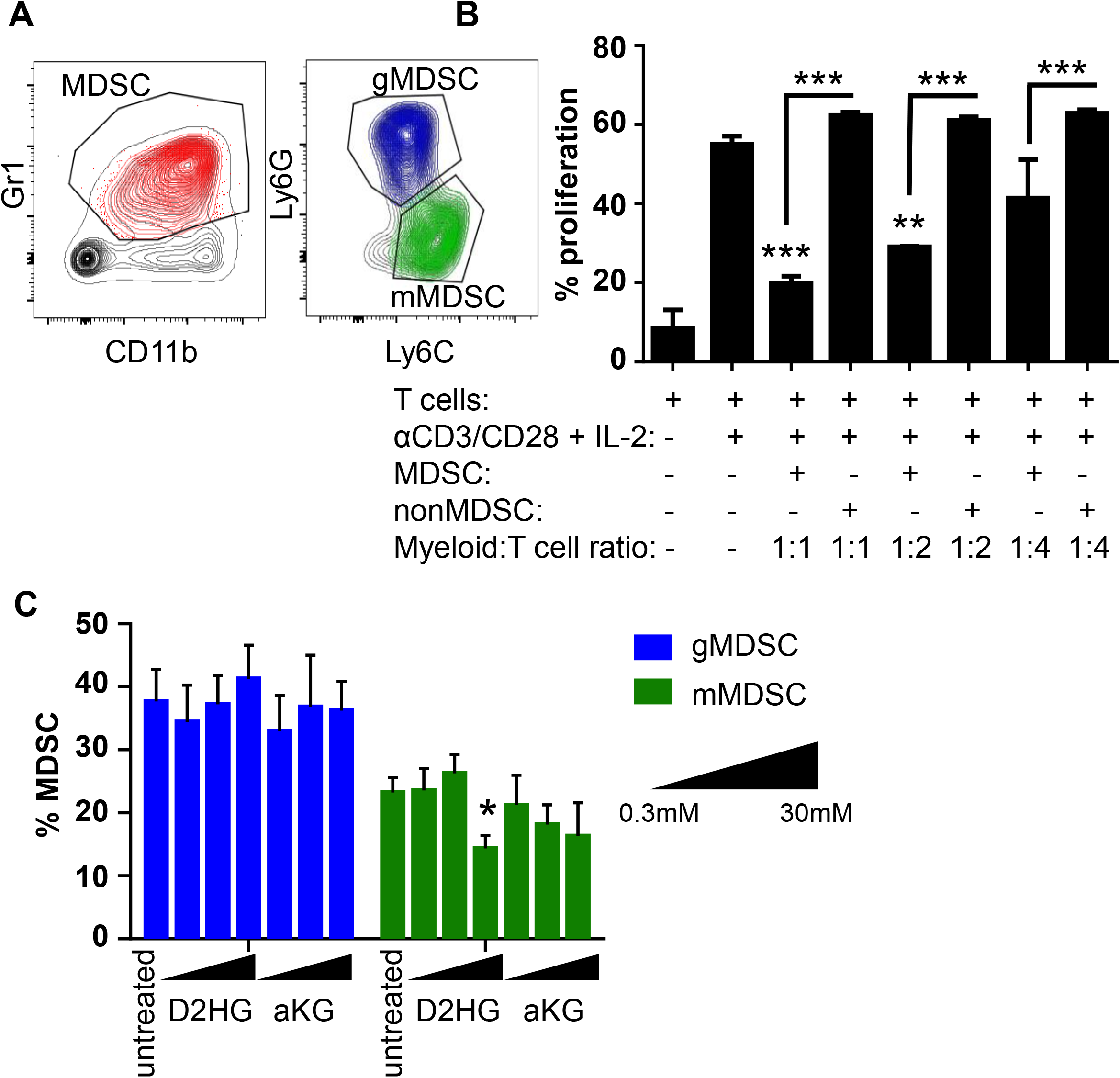
D2HG blocks mMDSC polarization in vitro. A) Representative flow plots demonstrating the gating strategy for in vitro generated mMDSCs and gMDSCs. B) Proliferation rate of T cells co-cultured with increasing concentrations of MDSCs or nonMDSCs. ** p<0.01 and *** p<0.001 based on unpaired t-test compared to the activated T cells. C) Percentage of gMDSCs and mMDSCs generated in the presence of αKG and D2HG. Data shown as mean ± SEM from n = 5. * p<0.05 based on unpaired t-test.

## Discussion

MDSCs have been linked to suppression of anti-tumor immune response and are emerging targets of cancer immunotherapy. Previous studies have demonstrated that the frequency of MDSCs in the peripheral blood is increased in various malignancies, including HCC^13–16,18^, and can impair therapeutic response^9,10^. Here, we demonstrated that different types of liver malignancies have distinct MDSC profiles. Despite increased MDSCs in blood of patients with HCC, CRLM and NET compared to benign lesions, MDSC levels were lower in CCA. CCA is broadly categorized as intrahepatic or extrahepatic, which vary in symptoms, prognosis and treatment response^25^. In our cohort, all but one patient with CCA had intrahepatic subtype preventing assessment of any potential correlations between CCA localization and MDSC accumulation pattern. IDH1/2 mutation in gliomas was shown to have a distinct immune infiltration profile compared to the wild type gliomas, represented as reduced leukocyte infiltration^26^. A recent study by Unruh et al., further demonstrated that although IDH mutation across tumor types lead to a unique methylation profiles, downregulation of immune-related genes is a common occurance^27^. Consistently, our studies established that the oncometabolite D2HG but not αKG inhibits MDSC polarization suggesting that increased D2HG abundance might be a conserved immune regulatory mechanism across IDH mutant tumors.

MDSCs subsets differ in their suppressive activity, ability to infiltrate tumors and disease associations. A recent study with a breast cancer model demonstrated that mMDSCs can modulate the local tumor microenvironment by interacting with cancer stem cells, while gMDSCs promote lung metastasis^19^. Consistent with the previous observations^14–16^, our results demonstrated that mMDSC subset (CD14^+^/CD15^-^/HLA-DR^-^) constitutes the majority of MDSCs detected in circulation of patients with primary or metastatic liver tumors. However, the frequency of gMDSCs can be affected by sample storage conditions^28^ and further analysis is required to determine whether the frequency of this population is augmented in hepatobiliary malignancies.

One of the MDSC-dependent immunosuppressive mechanisms is induction of Tregs^21^. Kalathil et al. demonstrated that compared to healthy controls the frequency of FoxP3^+^CTLA4^+^ Tregs was increased in HCC patients^13^. However, we did not observe a difference in the peripheral blood levels of CD25^+^CD127^-^ Tregs in patients with malignant liver tumors. While FoxP3 is widely used as a biomarker of Tregs, lack of CD127 expression more reliably marks activated Tregs^29^. Importantly, our patient cohort excludes any viral HCC cases. HBV/HCV infection has shown to recruit Tregs to restrain anti-viral immunity and inflammation-related liver damage^30,31^. Therefore, it is possible that viral HCC patients have more Tregs than non-viral HCC patients. While 75% of the HCC cases result from chronic HBV/HCV infection, non-viral HCC patients are diagnosed with higher tumor burden, pointing out the importance of considering HCC etiology^32^. ^6^. MDSCs also support tumorigenesis by attenuating cytotoxic T lymphocyte activity. By using CD107a expression as a marker of effector activity^33^, we investigated potential changes in activated T cells. As expected, the percentage of activated CD8^+^ T cells were relatively low in circulation but with no apparent difference between disease types.

Studies in breast cancer and glioblastoma have demonstrated that circulating MDSC frequency is elevated with tumor grade^34,35^. In our HCC cohort, two patients with the highest MDSC frequency had grade 3 tumors but due to limited number of grade 3 tumors further analysis could not be performed. There was no correlation between MDSC percentages and other clinical parameters such as TNM stage, AST, ALT or tumor markers levels. This indicated that MDSC frequency is an independent predictor of malignancy and not confounded by clinical variables in HCC. ^36^

Unlike gMDSCs, which are hypothesized to be terminally differentiated, mMDSCs can polarize into DCs and macrophages ^37,38^. Since mMDSCs were the prevalent cell type in patients with liver cancer, we speculated that mMDSC-APC differentiation axis was hindered leading to accumulation of the former population in tissues. Lower number of intralesional CD33 positive cells in malignant tumors pointed out to a potential reduction in myeloid infiltration. It is also possible that the CD33 staining intensity is biasing the results, as APCs express higher levels of CD33 compared to immature myeloid cells^39^. In contrast, the decreased number of HLA-DR positive cells in CCA and HCC with respect to the benign lesions may suggest a reduced ability to elicit an immune response in these tumors.

In line with this observation, HCC patients with high MDSC score (CD33^high^CD11b^high^HLA-DR^low^) in the TCGA dataset, had survival disadvantage. One possible explanation of decreased CD33 and HLA-DR positive cells in malignant tumors is that ENTPD2/CD39L expression by tumor cells hinders MDSC maturation in liver tumors^40^. Although this is yet to be confirmed in HCC patients, low MDSC frequencies in CCA patients suggest that there might be alternative mechanisms impairing trafficking or maturation of APCs in this tumor type.

Collectively, our results identify a distinct immune signature presented as high circulating MDSC levels and low tumor-infiltrating APCs in non-viral HCC as opposed to low peripheral MDSCs and low tumor-infiltrating APCs in CCA, and low circulating MDSCs and more intrahepatic APCs in benign lesions (**Fig 5**). Our findings make the case for further studies investigating a role of blood MDSC measurements as a diagnostic test for primary and metastatic hepatobiliary malignancies and potential role of therapeutic targeting. The use of metronomic chemotherapy for depletion of MDSCs^11,41–44^ is currently under clinical evaluation for glioblastoma and non-small cell lung cancer [NCT02669173, NCT03302247]. This approach compromises a promising therapeutic opportunity for HCC, CRLM and NETs as well.

**Figure 5.**
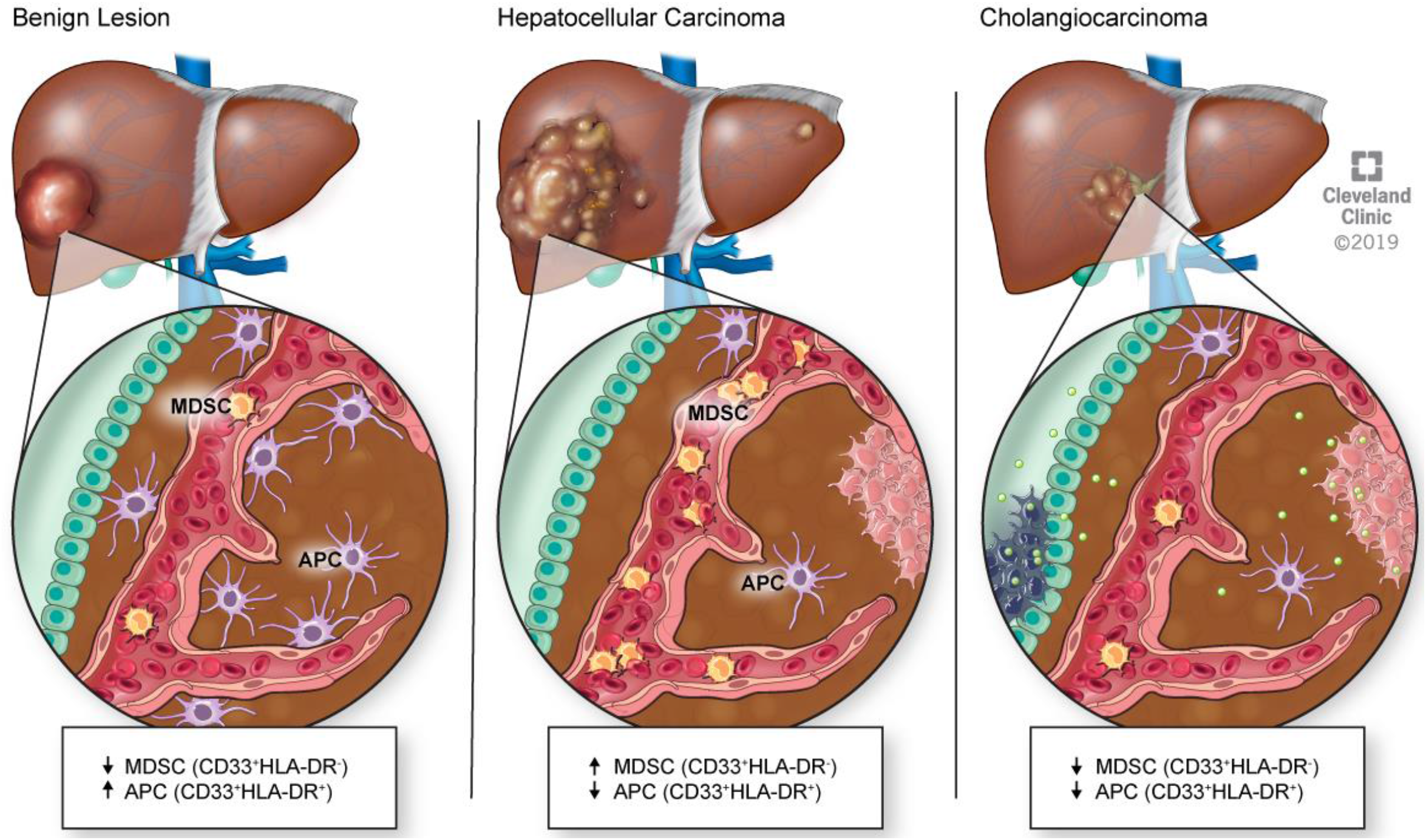
Malignant liver cancers have more circulating MDSCs and less tumor-infiltrating APCs.

## Acknowledgements

The authors would like to thank the members of the Lathia laboratory, Eric Shultz and Joseph Gerow from the Lerner Research Institute Flow Cytometry Core and Amanda Mendelsohn from the Center for Medical Art and Photography at the Cleveland Clinic for their assistance. We are also grateful to the patients at the Cleveland Clinic for consenting to the use of their tissue and blood for these studies.

## Abbreviations

AFP: Alpha-fetoprotein
αKGα: Alpha-ketoglutarate
AST: Aspartate aminotransferase
ALT: Alanine transaminase
APC: Antigen-presenting cells
CCA: Cholangiocarcinoma
CEA: Carcinoembryonic antigen
CRLM: Colorectal liver metastases
D2HG: D-2-Hydroxyglutarate
DC: Dendritic cell
gMDSC: Granulocytic myeloid-derived suppressor cells
HCC: Hepatocellular carcinoma
MDSC: Myeloid-derived suppressor cells
mMDSC: Monocytic myeloid-derived suppressor cells
NET: Neuroendocrine tumor
PBMC: Peripheral blood mononuclear cells
TCGA: The Cancer Genome Atlas
Treg: T regulatory cells
WBC: White blood cells

---

**Supplemental Figure 1.**
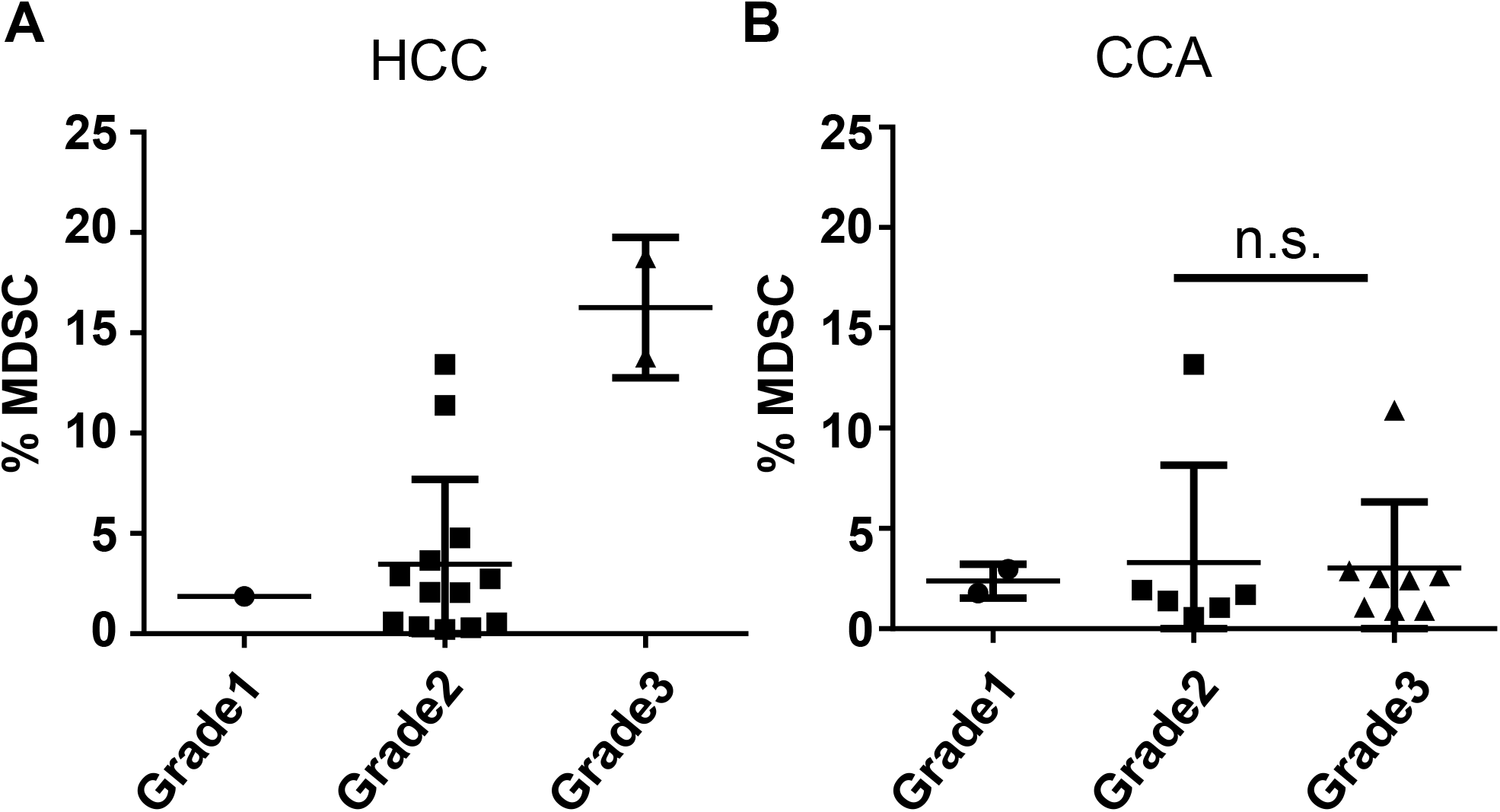
Low MDSC frequency is independent of grade of CCA. Percentage of MDSCs in A) HCC and B) CCA patients based on grade.

**Supplemental Figure 2.**
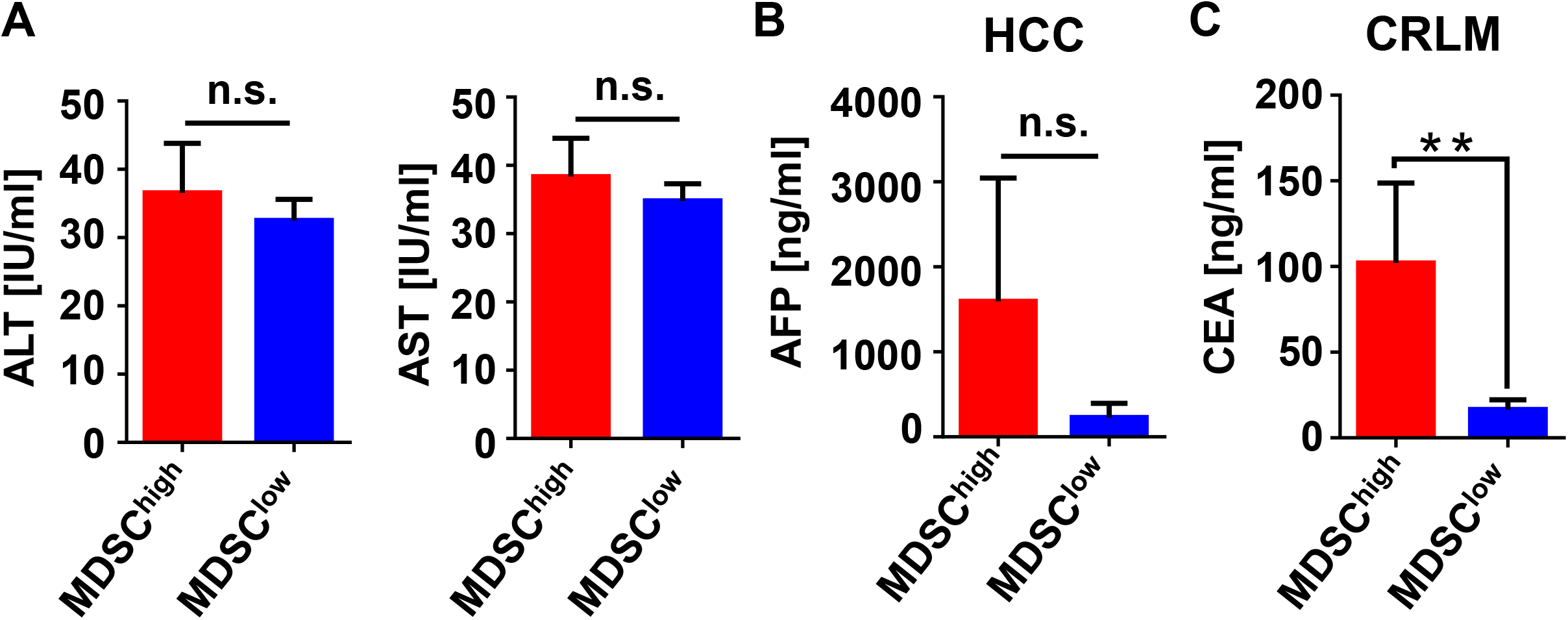
MDSC frequency in peripheral blood of HCC, CRLM and NET is independent of other clinical parameters. A) ALT and AST levels in all patients subdivided into MDSC high (>4.5%, n=23) versus low (<4.5%, n=86) group. B) AFP levels are similar in HCC patients with high (n=6) versus low (n=13) MDSC score. C) Increased CEA levels in CRLM patients with high (n=11) versus low (n=29) MDSC score ** p<.01 as determined by two-tailed t-test with equal variance.

**Supplemental Table 1:** Immunohistochemistry staining protocol

|  |  |  |  |
| --- | --- | --- | --- |
| **Antibody** | **CD33** | **HLA-DR** | **CD68** |
| **Clone** | PWS44 | CR3-43 | KP-1 |
| **Manufacturer** | Novocastra<br>(Leica) | ThermoFisher | Ventana |
| **Dilution** | 1:100 |  | Pre-dilute |
| **Auto-Stainer** | Ventana BenchMark Ultra |  |  |
| **Ab Incubation Time** | 40 minutes |  | 16 minutes |
| **Ab Incubation Temp** | 37 Deg C |  |  |
| **Retrieval Buffer** | Ventana CC1 |  |  |
| **Antigen Retrieval** | Select 64 minutes (STD) |  | Select 8 minutes |
| **Detection Kit** | Ventana iView<br>with A/B blocker | Ventana ultraView<br>with Mouse Ab AMP | OptiView DAB detection<br>(pre-primary peroxidase inhibitor selected) |
| **Positive Control Tissue** | Normal Lymph Node |  | Normal Tonsil |

